# Deep Learning models for retinal cell classification

**DOI:** 10.1101/2023.05.26.542384

**Authors:** Maciej Kostałkowski, Katarzyna Kordecka, Jagoda Płaczkiewicz, Anna Posłuszny, Andrzej Foik

**Author notes:** Corresponding author;, address: Kasprzaka 44/55, 01-224 Warsaw, Poland.

## Abstract

Data analysis is equally important as an experimental part of the scientist’s work. Therefore any reliable automatization would accelerate research. Histology is a good example, where scientists work with different cell types. The difficulty level can be severe while trying to distinguish cell types from one another. In this paper, we focus on the retina. The retina consists of eight basic cell types, creating a layered structure. Some types of cells overlap within the layer, and some differ significantly in size. Fast and thorough manual analysis of the cross-section is impossible. Even though Deep Learning models are applied in multiple domains, we observe little effort to automatize retinal analysis. Therefore, this research aims to create a model for classifying retinal cell types based on morphology in a cross-section of retinal cell images.

In this study, we propose a classification Deep Learning model for retinal cell classification. We implemented two models, each tested in three different approaches: Small dataset, Extended dataset, and One cell type vs. All cell types. Although the problem presented to the trained model was simplified, a significant data imbalance was created from multiclass to binary classification, influencing the models’ performance. Both, Sequential and Transfer Learning models performed best with the Extended dataset. The Sequential model generated the best overall results. The obtained results allow us to place prepared models within the benchmark of published models.

This paper proposes the first Deep Learning tool classifying retinal cell types based on a dataset prepared from publicly available images collated from multiple sources and images obtained in our laboratory. The multiclass approach with an extended dataset showed the best results. With more effort, the model could become an excellent analytical tool.

## 1. Introduction

Data analysis is one of the most time-consuming parts of a scientist’s work. Therefore, any reliable automatization would speed up research. The intensive growth of computer vision-based systems followed the interest of big technology companies in Deep Learning (DL) algorithms in 2015. Since then, many new tools have been and are developed. Such tools are crucial when analyzed data is complicated, and information is packed densely packed.

One of the domains that started to leverage DL algorithms heavily is histology. Some applications focus on cell classification in real time under the microscope (Gu et al., 2021), while others use the DL models on captured images (Coudray et al., 2018; Iqbal et al., 2021; Ker et al., 2019; Tabibu et al., 2019; Zhang et al., 2017). Most models perform binary classification, deciding whether the presented image shows healthy or diseased tissue fragments (Iqbal et al., 2021; Zhang et al., 2017). Due to minor differences between different cell structures, there is a relatively low number of publications applying a multiclass approach (Coudray et al., 2018; Gu et al., 2021). To simplify the classification task, Tabibu and coworkers used decision three consisting of binary classifiers, to compare the most characteristic classes against each other (Tabibu et al., 2019). The transfer learning technique is used whenever authors work with a small dataset that is hard to extend. Knowledge from the pre-trained model is applied to the presented task to boost the results. The Inception V3 architecture is a common choice for transfer learning (Coudray et al., 2018; Ker et al., 2019). Another approach to successfully improve results is to, use own well-working model to support the model with a poor performance instead of using a pretrained model from public resources (Zhang et al., 2017).

Despite different approaches, we still need solutions to automatize the analysis of microscope images based on cell morphology. Therefore, this research aims to develop a retinal cell type classification model based on cross-sections of retinal cell images. The model focuses on classifying eight basic retinal cells types, such as Amacrine Cell (AC), Bipolar Cell (BC), Cone, Retinal Ganglion Cells (RGC), Horizontal Cell (HC), Muller Cell (MC), Rod, and Retinal Pigment Epithelium (RPE). Additionally, we considered multiple scenarios related to different types of sequential models and dataset sizes to achieve the best possible efficiency.

In this paper, we trained two sequential and transfer learning models. We collected images of retina cells available in publications and our laboratory for a dataset and prepared an extended version. To simplify the problem, we tested the “One cell type vs. All cell types” approach. Despite the limited dataset, the best model was the sequential model with the extended dataset.

## 2. Methods

### 2.1 Images from the lab resources

The tissue used to capture retinal cell images was obtained from the animal study reviewed and approved by the First Warsaw Local Ethical Commission for Animal Experimentation (1194/2021).

To identify retinal cells, we use two strategies: immunofluorescence staining with antibodies and intravitreal injection of a specific viral vector. In both cases, we sacrifice mice with a lethal dose of pentobarbital. Eyes are enucleated, fixed with 4% paraformaldehyde for two hours at 4 °C, and then washed with PBS. After that, the eyes are moved into 5, 10, 20, and 30% of the sucrose solution until the eyeballs for cryoprotection. Such prepared tissue can be frozen in the OCT medium, cut to 15 µm cryosections, and mounted on glass slides.

For immunofluorescence staining, slices are incubated in 5 % normal bovine serum with 0.2% Triton X-100 in PBS for one h and incubated overnight with combinations of different primary antibodies diluted in PBS containing 1% Triton X-100 and 2% BSA. Primary antibodies were selected for each cell type. In our studies, we are going to use the following antibodies to stain specific cell types: BRN3A for Ganglion cell types (Krieger et al., 2017), PNA for cones (Krishnamoorthy et al., 2008), rhodopsin for rods (Gargini et al., 2017), CALB for horizontal cells (Cuenca et al., 2014), PKC-_α_ for bipolar cells (Strettoi et al., 2004) and GAD67, PV, and ChAT for Amacrine cells (de Sevilla Müller et al., 2017), GFAP for Muller glia (Ren et al., 2018), RPE65 for RPE (Gamm et al., 2008). Subsequently, the sections were washed in PBS and incubated in the secondary antibodies for one h. The sections were finally washed in PBS and mounted in DABCO medium.

Intravitreal injection of viruses: The intravitreal injection of 1 µl viral solution will be performed into both eyes. After a few weeks (3-4), the animals were sacrificed. Eyes will be cut in the cryostat for retinal sectioning, as described above.

Images captured with the highest possible resolution will be stored on the PC. Before their usage, each image will have to be marked manually. Processing will include finding and scoping a proper cell example in the photo.

### 2.2 Models

Due to its performance, convolutional Neural Network (CNN) is the most common DL model type used for object classification. A typical CNN model consists of convolutions creating a high-level feature representation of an image, followed by the neuronal network. The network consists of layers created by the nodes. Each node is a value established during the training process. During detection, the result is estimated by verifying which set of nodes associated with the class was the most activated.

First, we implemented a simple sequential model, figure 1(a). The model consists of three layers of image transformation involving data augmentation using random rotation and random zoom and ending with rescaling. Next are three steps of convolutions and max-pooling for feature extraction and information reduction, respectively. Each convolution layer has a kernel set to 3, the same padding, and a hidden rectifier linear unit implemented. Then the image is fed forward to the dropout layer, which randomly rejects half of the extracted features to reduce the overfitting effect. The remaining image features are transformed into a flattened layer and forwarded to two fully dense perceptron layers ending with a node for each class. The first dense layer has additionally implemented L2 regularization with the setting of 0.01 with a hidden rectifier linear unit included. The remaining details of the sequential model are presented in the table, figure 1(a).

**Figure 1.**
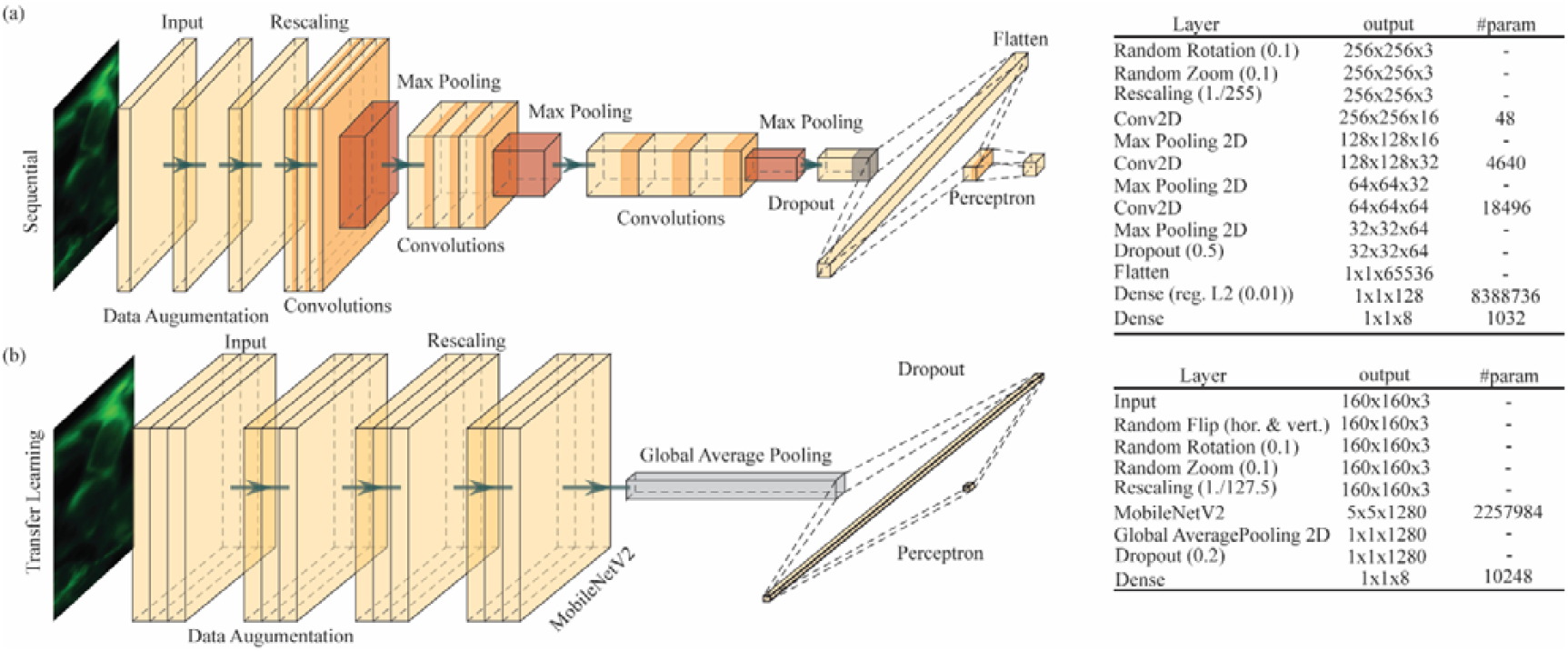
Visualizations of Convolutional Neural Network models used in this study. **(a)** The sequential model consists of three layers of image formation followed by three convolution operations with max pooling, a dropout layer, flatten layer, and two layers of a fully dense perceptron. **(b)** The transfer learning model consists of image transformation layers, and a global pooling layer merging weights from the MobileNetV2 model (Sandler et al., 2018) with information from the input image. The model ends with two layers of a fully dense perceptron.

The second model we used implements transfer learning, figure 1(b). The model starts with three layers of data augmentation: random flip, random image zoom, and random rotation. Next, the image goes through the MobileNetV2 model (pretrained network) (Sandler et al., 2018), followed by a global pooling layer to flatten the output. The model ends with a two-layer perceptron consisting of a dropout layer and a dense layer with nodes equal to the number of classes. Detailed layer settings are provided in the figure 1(b) table.

### 2.3 Metrics

We introduced basic parameters such as Accuracy and Loss to understand and quantify a model’s performance. Accuracy describes the ratio of positive recognitions and overall occurrences. The Loss is a cross-entropy function describing the distribution q relative to a distribution p over the given set:

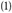

where Ep[.] is the expected value operator with respect to the distribution p.

Both parameters are often recorded during training and validation, and displayed in the plot, allowing us to observe whether the overfitting or underfitting effect occurs. The goal is to achieve as high as possible Accuracy while keeping Loss as low as possible.

Depending on the aim of the model, parameters may vary. Four counters need to be recorded to capture the most important parameters to validate our model. The True Positive (TP), to see how many objects are correctly recognized; the True Negative (TN) to see how many objects are correctly rejected; the False Positive (FP) how many objects are incorrectly recognized; and the False Negative (FN) to how many are incorrectly rejected. The Accuracy can be calculated using the formula:

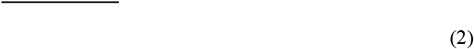

Although this can be very insightful information about the model, it cannot predict its performance. Thus, we implemented additional metrics:

Precision - to describe the specificity of a model,

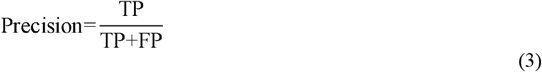

Recall - to describe the selectivity of a model,

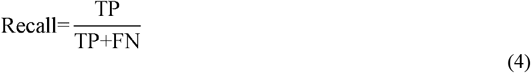

F1 score - to describe the balance between both previous parameters.

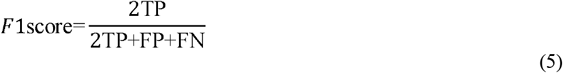

### 2.4 Dataset preparation

The dataset we prepared is based on photos and schematics of a cross-section retina publicly available online or from scientific publications (Boije et al., 2016; Esquiva et al., 2017; Fain & Sampath, 2018; Grimes et al., 2021; Jusuf & Harris, 2009; Masland, 2001; Poché & Reese, 2009; Wan & Goldman, 2017; Wilson & Di Polo, 2012; Zele & Cao, 2014). To satisfy classifier needs, we have cropped out individual cells from the collected images (∼35), figure 2(a-h), and assigned them to an appropriate folder, figure 2(i). We the size of the cropping window, we have adjusted to each object individually. The single image should contain a body cell and a visible part of the dendrites associated with the targeted body cell.

**Figure 2.**
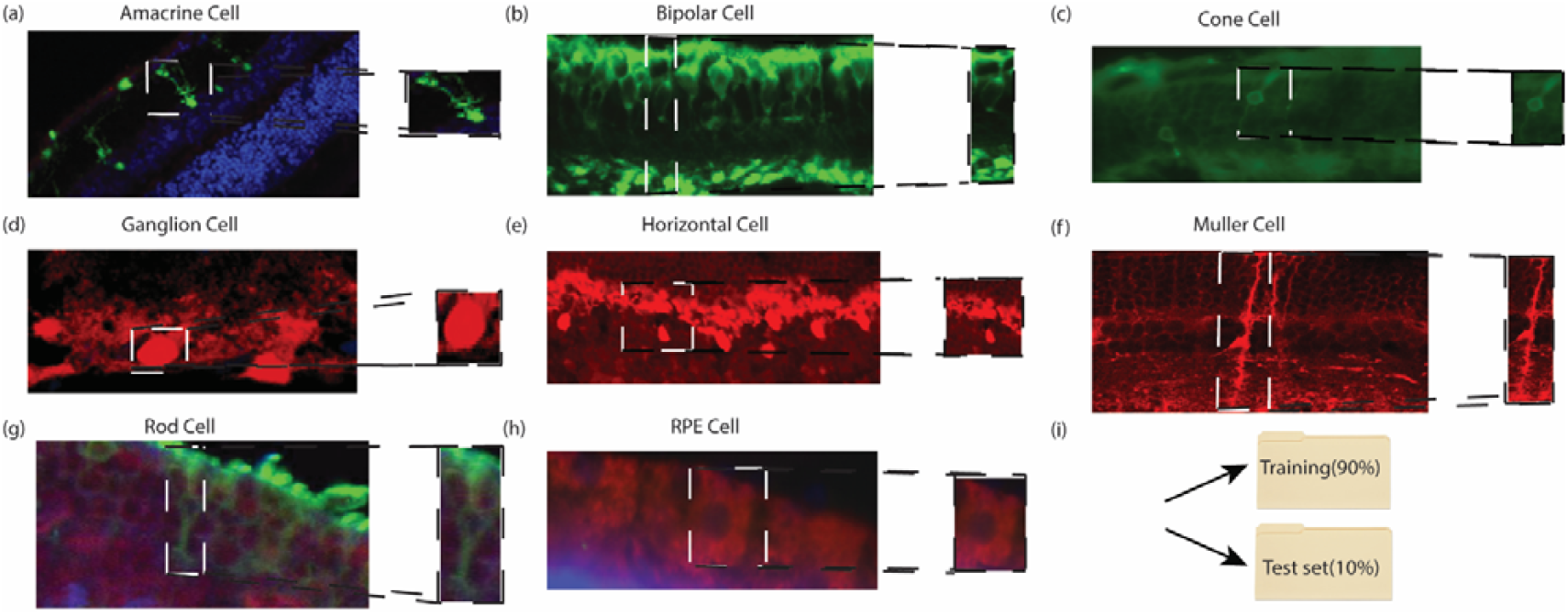
Process of the dataset preparation. An image of a selected cell was cropped out from the cross-section retina image. The size of the cropping window was adjusted for each object individually. The image should contain a body cell and a visible part of the dendrites associated with the targeted body cell. Images were captured with 20x magnification. **(a)** Example of an Amacrine Cell in a cross-section of the retina. **(b)** Example of a Bipolar cell in a cross-section of the retina. **(c)** Example of a Cone cell in a cross-section of the retina. **(d)** Example of a Retinal Ganglion Cell in a cross-section of the retina. **(e)** Example of a Horizontal cell in a cross-section of the retina. **(f)** Example of a Muller cell in a cross-section of the retina. **(g)** Example of Rod cell in a cross-section of the retina. **(h)** Example of an RPE cell in a cross-section of the retina. **(i)** Cropped image is saved in the appropriate folder gathering the same cell types. 90% of the collected images are dedicated to training and validation. The remaining 10% is stored in a separate folder for testing purposes.

Initially, we started with 96 Amacrine Cells (AC), 104 Bipolar Cells (BC), 74 Cone cells, 102 Retinal Ganglion Cells (RGC), 33 Horizontal Cells (HC), 63 Muller Cells (MC), 81 Rod cells, and 76 Retinal Pigment Epithelium (RPE) cells. In total, the size of the dataset was 629 images of the different samples. Before training, the dataset we split into 90%/10% for training and testing. The code used for both models automatically divides the training part of the images and splits it into 80%/20% proportion for training and validation. Thus, the final data split was 72%/18%/10%, with 446/98/85 examples. After initial training and testing, the dataset was increased in GIMP software by applying grayscale, noise addition (0.3 random factor setting for each channel), and edge extraction (gradient algorithm, 5.0-factor setting, no behavior for edges) macros. Thus the dataset ended at 2960 examples. The exact numbers describing small and extended datasets are shown in Table 1.

**Table 1.** Dataset preparation for each cell type.

| Cell type | Number of cells | Training/ Validation | Test |
| --- | --- | --- | --- |
| Amacrine | 96(456) | 84(420) | 12(36) |
| Bipolar | 104(488) | 87(452) | 17(36) |
| Cone | 74(360) | 63(328) | 11(32) |
| Ganglion | 102(408) | 95(380) | 7(28) |
| Horizontal | 33(192) | 28(160) | 5(32) |
| Muller | 63(316) | 53(284) | 10(32) |
| Rod | 81(380) | 68(352) | 13(28) |
| RPE | 76(360) | 66(332) | 10(28) |
| Sum | 629(2960) | 544(2708) | 85(252) |

Numbers show numbers for the small dataset and extended dataset in the brackets.

### 2.5 Training procedure

We trained Sequential models with Stochastic Gradient Descent (SGD) optimizer, the learning rate 0.01, and Nestorov momentum activated. The loss function uses sparse categorical cross entropy calculated directly from logits.

For Transfer learning models, we selected almost identical settings. The only changes are in a lower learning rate of 0.001 in the initial training phase and 0.0001 at the fine-tuning phase, and momentum set to 0.9.

Additionally, we implemented early stopping for both types of models to monitor the loss function of the validation dataset with a minimum required improvement set for 0.01 and patience of 10 epochs.

### 2.5 Testing procedure

10% of the cell images we collected, were saved for testing. We applied the test set to a pre-trained network and collected TP, TN, FP, and FN counters based on model predictions. This allowed us to calculate all basic metrics to describe the model’s performance. Additionally, we prepared Region Operating Curve (ROC) plots for graphical performance representation. To average the models’ performance, we prepared a final evaluation based on results gathered from testing repeated 10 times.

### 2.6 Equipment

All models were tested and validated on Intel Core i7-10700 CPU 2.9GHz, 16GB, NVIDIA GeForce RTX 2070 SUPER using CUDA 11.4 toolkit.

## 3. Results

To obtain our results, we have prepared two models, a sequential model and a model using transfer learning. To quantify objectively models’ performance, we used the following metrics: Precision, Recall, F1, and Accuracy. Additionally, we plot ROC for each model. The dataset images required for the training were partially found in available publications and partially collected in our laboratory. We started training both models with 629 samples. Then we repeated the process with an artificially extended dataset (4x). We have also tested the “One cell type vs. All cell types” (One vs. All) approach. Ultimately, we present how models work with previously unseen cell examples.

### 3.1 Model training using a small dataset

Initially, we trained a sequential model with 629 samples using the early stopping technique to prevent the overtraining effects. We set a monitoring function to watch over the value loss parameter with patience of 10 epochs. We achieved the best results using Stochastic Gradient Descent (SGD) optimizer with the Nesterov momentum enabled and a learning rate set at 0.01. Training results for each class are shown in Table 2 and plotted ROCs (Figure 3(a)). The Accuracy column shows a rating of over 86%. However, an in-depth analysis reveals the best Precision for both, Amacrine cells (AC) and Muller Glia (MC) classes and the worst for Retinal Ganglion Cells (RGC). In the Recall category, the Rod class reached perfect scores of 100% and the worst, 14%, for the ganglion class. The F1 score shows that the most balanced class is MC, 94%, followed by 22% for RGC. ROC plot similarly reveals the worst performance for RGC class. We obtained the best results for the Rods class, 86% in Precision and 100% in Recall.

**Table 2.** Table showing detailed metrics for both models with Small, Extended dataset and the One vs. All approach.

|  |  | Multiclass<br>Small Dataset |  |  |  | Multiclass<br>Extended Dataset |  |  |  | Binary<br>One vs All |  |  |  |
| --- | --- | --- | --- | --- | --- | --- | --- | --- | --- | --- | --- | --- | --- |
|  |  | Precision | Recall | F1 | Accuracy | Precision | Recall | F1 | Accuracy | Precision | Recall | F1 | Accuracy |
| Sequential | Amacrine | 1.0 | 0.84 | 0.91 | 0.98 | 1.0 | **0.96** | **0.98** | **0.99** | 0.79 | **0.96** | 0.87 | 0.96 |
|  | Bipolar | 0.61 | 0.83 | 0.7 | 0.86 | **1.0** | **0.96** | **0.98** | **0.99** | 0.34 | 0.08 | 0.13 | 0.84 |
|  | Cone | 0.68 | 0.63 | 0.65 | 0.91 | **0.97** | **1.0** | **0.98** | **1.0** | **0.69** | **1.0** | **0.82** | **0.95** |
|  | Ganglion | 0.56 | 0.14 | 0.22 | 0.93 | **1.0** | **0.9** | **0.95** | **0.99** | 0.0 | 0.0 | 0.0 | 0.89 |
|  | Horizontal | 0.76 | 0.45 | 0.57 | 0.96 | **1.0** | **1.0** | **1.0** | **1.0** | **1.0** | 0.06 | 0.11 | 0.88 |
|  | Muller | 1.0 | 0.88 | 0.94 | 0.99 | 0.82 | **1.0** | 0.9 | 0.97 | 0.0 | 0.0 | 0.0 | 0.72 |
|  | Rod | 0.86 | 1.0 | 0.92 | 0.97 | **1.0** | 1.0 | **1.0** | **1.0** | 0.0 | 0.0 | 0.0 | 0.82 |
|  | Rpe | 0.74 | 0.91 | 0.82 | 0.95 | **0.99** | 0.91 | **0.95** | **0.99** | 0.32 | 0.21 | 0.25 | 0.81 |
|  |  | Precision | Recall | F1 | Accuracy | Precision | Recall | F1 | Accuracy | Precision | Recall | F1 | Accuracy |
| Transfer Learning | Amacrine | 0.87 | 0.85 | 0.86 | 0.97 | **1.0** | **0.9** | **0.95** | **0.99** | 0.38 | 0.2 | 0.26 | 0.82 |
|  | Bipolar | 0.86 | 1.0 | 0.92 | 0.97 | 0.81 | 0.98 | **0.89** | 0.97 | 0.0 | 0.0 | 0.0 | 0.84 |
|  | Cone | 0.66 | 0.81 | 0.73 | 0.92 | **1.0** | 0.79 | **0.88** | **0.97** | **0.91** | 0.8 | **0.85** | **0.96** |
|  | Ganglion | 0.86 | 0.82 | 0.84 | 0.97 | **1.0** | 0.78 | **0.88** | **0.98** | 0.0 | 0.0 | 0.0 | 0.89 |
|  | Horizontal | 1.0 | 0.78 | 0.88 | 0.99 | 1.0 | **1.0** | **1.0** | **1.0** | 0.0 | 0.0 | 0.0 | 0.88 |
|  | Muller | 1.0 | 0.55 | 0.71 | 0.94 | **0.72** | **1.0** | **0.84** | **0.95** | 0.29 | **1.0** | 0.45 | 0.71 |
|  | Rod | 0.8 | 1.0 | 0.89 | 0.96 | **1.0** | 0.92 | **0.96** | **0.99** | 0.15 | 0.2 | 0.17 | 0.78 |
|  | Rpe | 1.0 | 0.77 | 0.87 | 0.97 | 1.0 | **1.0** | **1.0** | **1.0** | 0.44 | 0.71 | 0.54 | 0.87 |

**Figure 3.**
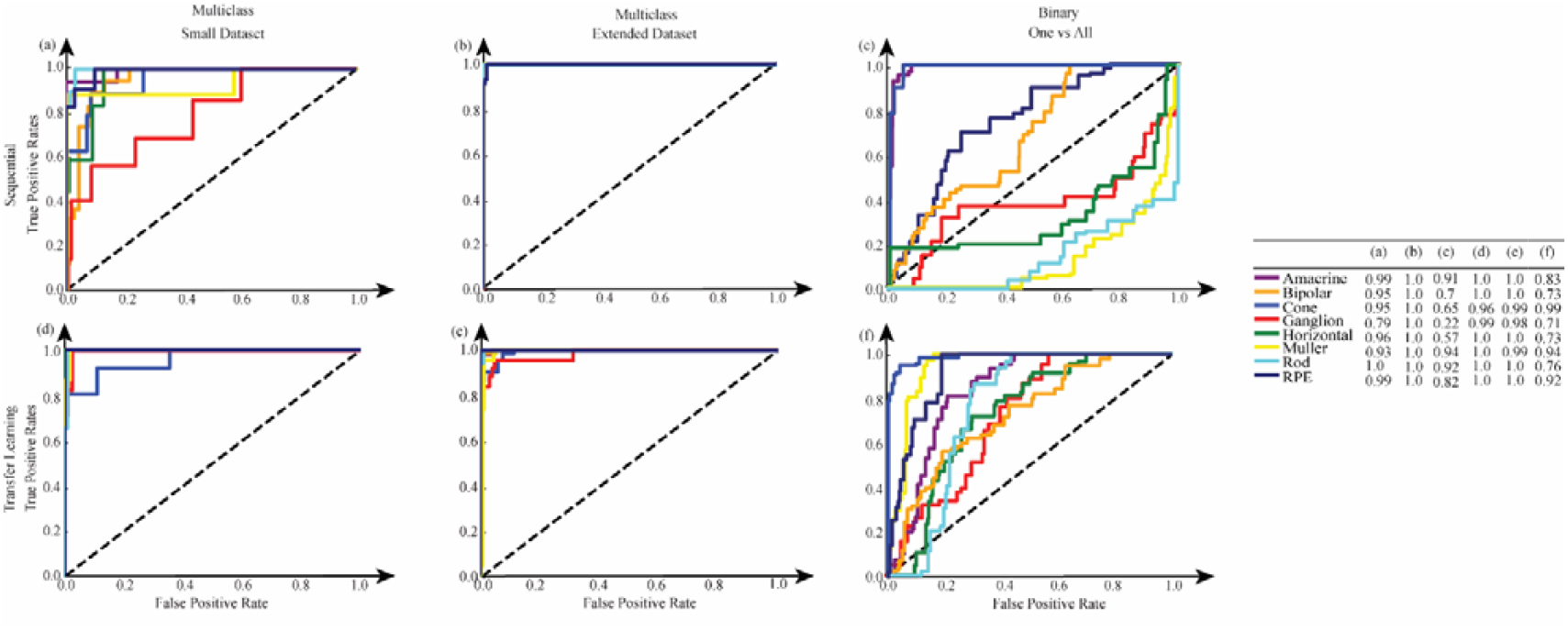
Region operating curves for the trained models. Each class has a color used for each plot. The curve illustrates the probability between True Positive versus False Positive image prediction. The Area Under the Curve (AUC) parameter expresses the curve in a number. Legend and AUC values for each plot are in the table on the right side. **(a)** the ROCs for a sequential model trained with a small dataset. **(b)** the ROCs for a sequential model trained with an extended dataset. **(c)** the ROCs for the One vs. All approach using sequential models. For each class, a separate model was trained using an extended dataset. **(d)** the ROCs for the transfer learning model trained with a small dataset. **(e)** the ROCs for the transfer learning model trained with an extended dataset. **(f)** the ROCs for the One vs. All approach using transfer learning models. For each class, a separate model was trained using an extended dataset.

In the next step, we have trained the transfer learning model. Similarly, we applied the early stopping technique to prevent the overtraining effect in the sequential model. We set the monitoring function to watch over the value of the loss parameter with patience of 10 epochs. We used the SGD optimizer with a momentum of 0.9 and a learning rate of 0.001, followed by a fine-tuning technique. In the last 100 layers of the MobileNetV2 model (Sandler et al., 2018), we trained with previously pre- trained layers using the same optimizer and momentum in the initial training phase. We decreased the learning rate to 0.0001 in both accuracy and loss function to prevent significant value changes and loss of information. Accuracy results for different classes span between the 92%-99% range, table 2. The maximum Precision value was reached in three classes, Horizontal Cell, MC, and RPE, with four scoring very high, AC, Bipolar Cell, RGC, and Rod cell, in the region of 80%-87%. The Cone class had adverse worse performance reaching 66% of Precision. The Recall metric has more varied results. Two classes reached perfect scores, and the worse performance was MC, with 55%. The MC class reached the lowest F1 score of 71% due to its weak Recall result. The other classes reached results from 73% to 92%. The ROC plot predicts the worst performance for the cone class and the perfect score for six classes, figure 3(d).

The first training round showed better performance of the transfer learning model overall. Even the worst classified class achieved better results than the sequential model, Table 2, Figure 3(a) and 3(d).

### 3.2 Model training using the extended dataset

Two suggestions were found in the literature while looking for methods to improve our models’ performance (Goodfellow et al., 2016). If possible, increase the dataset or simplify the task by changing the problem from multiclass to binary, known as the One vs. All approach. Since we had limited resources available over the internet, we extended the dataset by transforming existing files using three macros from the GIMP software as described in the Dataset Preparation section.

Applying an extended dataset, we repeated the training of sequential and transfer learning models. We have used the same training parameters as previously. For the sequential model, improvements are substantial. A score of 82%-100% was obtained for most of the cell classes. All metrics results are presented in Table 2 and Figure 3(b). The transfer learning model also improved, showing slightly worse metrics than the sequential model (72%-100%, Table 2 and Figure 3(e)).

Next, we tested the One vs. All approach. We trained both models to recognize each cell type separately against the remaining cell classes. For the sequential model, two classes scored above 90% in Accuracy, but RCG, MC, and Rod cells scored 0% in the Precision and the Recall metrics, Table 2 and Figure 3(c). Transfer learning models scored similarly. One class reached 90% and above Accuracy. Three classes scored 0% in Precision and Recall, BC, RGC, and HC. Additionally, AC and Rod cell types reached no more than 40% in Precision and Recall metrics, Table 2 and Figure 3(f).

### 3.3 Models use case

Lastly, we tested models with images not included in the training or testing dataset. We selected a representative number of images of each cell class and tested them on six pre-trained models, Figure 4. Models provided classification prediction and confidence estimation expressed in percentages for each image.

**Figure 4.**
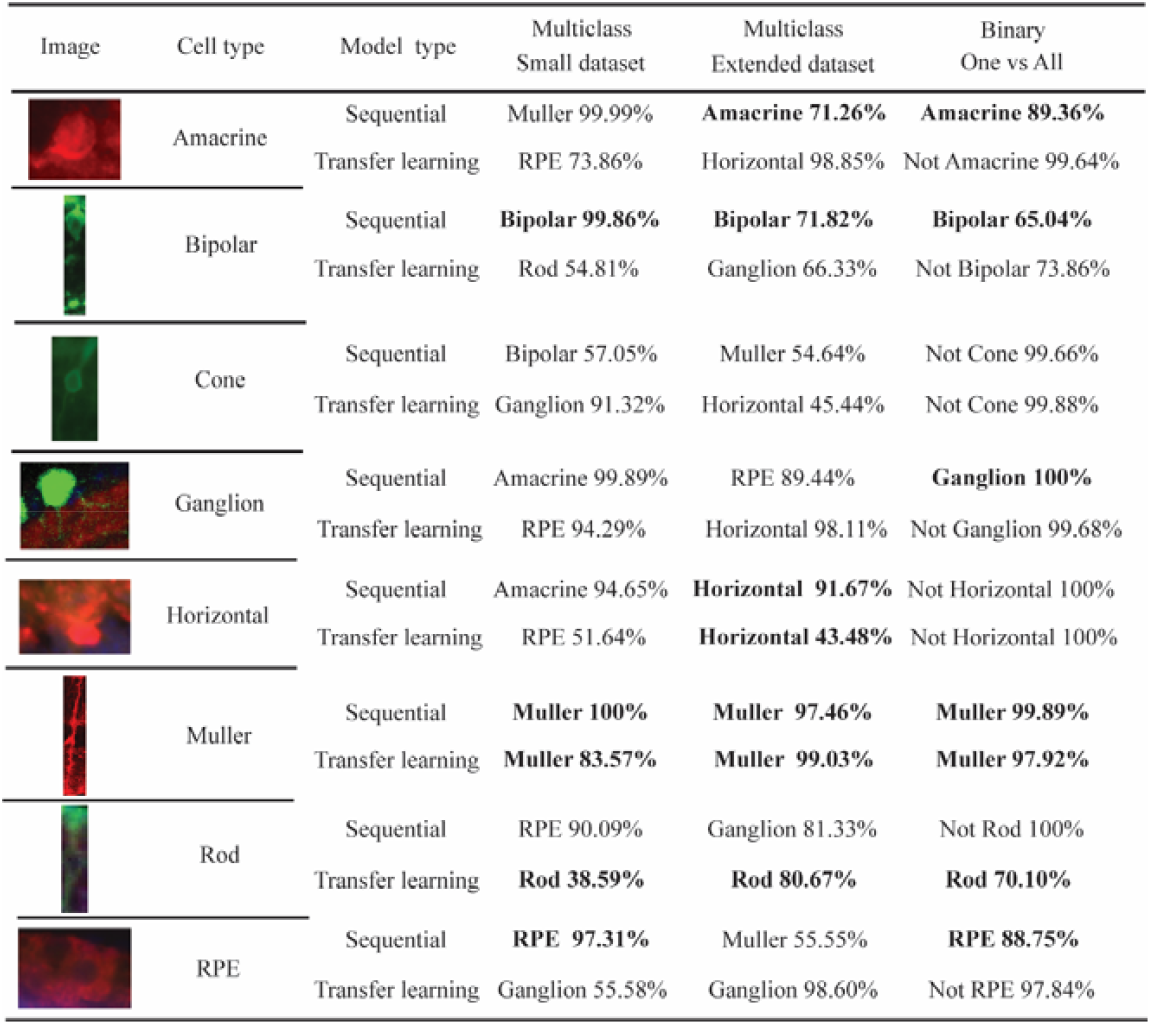
Example results with cells previously not seen by the models. Each cell was tested with each model. The results show predicted cell type names and prediction certainty expressed in percentages. Correct results are bolded.

MC had the best results; each model correctly classified the class. The cone cell example was the worst, and the example was not classified correctly at all. BC and Rod’s cells were classified correctly three times out of 6. AC and RPE cells twice, and RGC once. In total, models correctly classified 19 examples out of 36.

## 4. Discussion

We have proposed a retinal cell-type image classification method using the DL models for cell-type recognition. We implemented two models, and each was tested in three different approaches: Small dataset, Extended dataset, and One vs. All. The One vs. All approach did not perform as well as predicted. Although the problem presented to the trained model was simplified, a significant data imbalance was created from multiclass to binary classification. Even compensation techniques like under-sampling of all classes did not help to improve the results. Therefore, influencing the models’ performance. Sequential and Transfer Learning (TL) models improved when the Extended dataset was used. Initially, the Transfer Learning model scored better than the Sequential model for the Small dataset. The difference can be attributed to the influence of the TL weights from the MobileNetV2 model (Goodfellow lan, Bengio Yoshua, 2016). The Sequential model achieved the best overall results.

Similarly, as reported in the literature, the dataset was the biggest challenge (Noorbakhsh et al., 2020; Saha et al., 2019; Ting et al., 2019). Publicly available data related to the retina is very limited compared to the publicly available datasets related to other topics (Khan et al., 2021; Ting et al., 2019). Without a broad spectrum of examples, models will not have enough reliability to be used in the machine learning research practice despite reaching high training results. We tested different augmentation techniques and image transformations to improve the model’s performance. That certainly helps with a generalization of the model; however, it is still not an optimal solution (Goodfellow et al., 2016; He et al., 2021; Liu et al., 2019; Shorten & Khoshgoftaar, 2019). This project dataset consists of schematics and images of retinal cross-section fragments. The images included in our dataset are chosen for publication, the best from all the obtained in the experiment, which does not cover a typical research situation. Therefore, in the following stages of model development, the dataset needs to be extended with more typical examples.

A significant limitation of a classification model is that it will classify only one object at a time. The model will only classify the object with the highest probability in the case of multiple objects. Therefore, before the model can start classification, a user should crop out all objects of interest individually from the image. The size of the cropping window influences the models’ performance. Too big or small boundaries around the object can easily falsify the result. With multiple objects to process, cropping each of them separately and transferring between programs becomes a tedious process on its own. Therefore, it would be more efficient to use DL models that can process the entire image simultaneously, such as image segmentation models.

It is hard to compare DL models to each other if they are not trained on the same dataset and verified objectively. To make a valid comparison of our models, we selected three publications focusing on multiclass cell classification using DL models. Oei and coworkers proposed CNN based classification model for detecting intracellular actin networks (Oei Ronald Wihal AND Hou, 2019). They were using immuno-stained cells photographed under a microscope. The proposed model was based on VGG16 architecture designed for classification between 3 classes. The dataset consisted of about 500-600 samples per class. The used optimizer was Adam, with a batch size of 4, and the model was trained for 1000 epochs. The dataset split was unusual, 10:1 in proportion. Oei and coworkers employed transfer learning from ImageNet model and data augmentation to improve the results. In the result, they have achieved Accuracy between 93-97%. Shibata and coworkers prepared a multiclass cell classification model based on lectin microarray data (Shibata et al., 2020). Their dataset consisted of 1557 samples of human cells, divided into five classes. Four classes contain around 300-500 examples, and one class has 48 samples. The best proposed neural network model consisted of 2 hidden layers, 300 nodes each. The model was trained with the Adam optimizer and a low dropout of 0.1 for 120 epochs. As a result, the proposed models scored between 95.6%-98.6% in Accuracy. Shifat-E-Rabbi and coworkers trained six CNN-based models (Shifat-E-Rabbi et al., 2020). They have used four publicly available datasets to train models. The dataset consisted of HeLa cells divided into ten classes, human osteosarcoma cells divided into two classes, thyroid nuclei cells divided into three classes, and human epithelial cells divided into six classes. The best results were achieved by VGG16 based model and the Inception V3 model. Both models were improved with transfer learning with weights from the ImageNet model and data augmentations.

Shifat-E-Rabbi and coworkers tested the influence of transfer learning and data augmentations on the Accuracy. 5 out of 8 models improved their Accuracy by between 1.2%-15%. Later on adding data augmentation the results improved only in 4 cases by no more than 4%. The models scored between 62.4%-95.5% of Accuracy. In this paper we have proposed Sequential model and Transfer Learning models for retinal cell classification. The same as Shifat-E-Rabbi and coworkers we have used Deep Learning models instead of shallow networks as Oei and coworkers and Shibata and coworkers proposed. In our case classification was performed between 8 classes. In the beginning with 629 samples, model using transfer learning approach had better results than sequential model, however with more augmented dataset consisting of 2960 samples, sequential model achieved better results than transfer learning model. Models that we have presented in this study, scored between 86%-100% for each class in Accuracy.

## 5. Conclusions

This paper proposes the first DL tool for the classification of retinal cell types based on the dataset prepared from publicly available sources and images collected in the laboratory. The multiclass approach with an extended dataset showed the best results. With more effort, the model has the potential to become a good analytical tool in the future. Therefore, further DL model development is a part of the ongoing lab research.

## Data availability

Images used for the creation of the dataset were obtained from publicly available resources over the internet (Boije et al., 2016; Esquiva et al., 2017; Fain & Sampath, 2018; Grimes et al., 2021; Jusuf & Harris, 2009; Masland, 2001; Poché & Reese, 2009; Wan & Goldman, 2017; Wilson & Di Polo, 2012; Zele & Cao, 2014), including the following websites: retinalmicroscopy.com, clinicalgate.com/retina, webvision.med.utah.edu, neupsykey.com/the-visual-system-5/, commons.wikimedia.org/wiki/File:Ganglion_cell.svg, clinicalgate.com/retina/ and in the International Centre for Translational Eye Research, Institute of Physical Chemistry, PAS.

## Code availability

Models were prepared using tutorials available on the Tensorflow/Keras 2.0 website. Models were implemented with Python 3.9.6 edition using pyplot, NumPy, DateTime, io, itertools, sklearn, pandas, scikit, statistics, and scipy libraries. The codes are available to download from the GitHub repository (github.com/OBi-ICTER/DL-retina-cell-classifier). Visualization of the neuronal network was done with the ann visualizer 2.5 library.

## Acknowledgments

We want to thank Prof. Marinco Sarunic, Ph.D., for his helpful comments at the early stage of this manuscript preparation. The International Centre for Translational Eye Research (MAB/2019/12) project is carried out within the International Research Agendas program of the Foundation for Polish Science co-financed by the European Union under the European Regional Development Fund and the Sonata Bis 9 grant (2019/34/E/NZ5/00434) funded by National Science Center for ATF.

## Declaration of generative AI in scientific writing

The authors declare that no generative artificial intelligence(AI) and AI-assisted technologies have been used in the writing process in this paper.

## References

Boije, H., Shirazi Fard, S., & Edqvist Per-Henrikand Hallböök, F. (2016). Horizontal cells, the odd ones out in the retina, give insightsinto development and disease. Front. Neuroanat., 10, 77.

Coudray, N., Ocampo, P. S., Sakellaropoulos, T., Narula, N., Snuderl, M., Fenyö, D., Moreira, A. L., Razavian, N., & Tsirigos, A. (2018). Classification and mutation prediction from non–small cell lung cancer histopathology images using deep learning. Nature Medicine, 24(10), 1559–1567. https://doi.org/10.1038/s41591-018-0177-5

Cuenca, N., Fernández-Sánchez, L., Sauvé, Y., Segura, F. J., Martínez-Navarrete, G., Tamarit, J. M., Fuentes-Broto, L., Sanchez-Cano, A., & Pinilla, I. (2014). Correlation between SD-OCT, immunocytochemistry and functional findings in an animal model of retinal degeneration. Frontiers in Neuroanatomy, 8. https://doi.org/10.3389/fnana.2014.00151

de Sevilla Müller, L., Azar, S. S., de los Santos, J., & Brecha, N. C. (2017). Prox1 Is a Marker for AII Amacrine Cells in the Mouse Retina. Frontiers in Neuroanatomy, 11. https://doi.org/10.3389/fnana.2017.00039

Esquiva, G., Lax, P., Pérez-Santonja, J. J., Garc\’\ia-Fernández, J. M., & Cuenca, N. (2017). Loss of melanopsin-expressing ganglion cell subtypes anddendritic degeneration in the aging human retina. Front. Aging Neurosci., 9, 79.

Fain, G., & Sampath, A. P. (2018). Rod and cone interactions in the retina. F1000Res., 7.

Gamm, D. M., Melvan, J. N., Shearer, R. L., Pinilla, I., Sabat, G., Svendsen, C. N., & Wright, L. S. (2008). A Novel Serum-Free Method for Culturing Human Prenatal Retinal Pigment Epithelial Cells. Investigative Ophthalmology & Visual Science, 49(2), 788–799. https://doi.org/10.1167/iovs.07-0777

Gargini, C., Novelli, E., Piano, I., Biagioni, M., & Strettoi, E. (2017). Pattern of retinal morphological and functional decay in a light-inducible, rhodopsin mutant mouse. Scientific Reports, 7(1), 5730. https://doi.org/10.1038/s41598-017-06045-x

Goodfellow, I., Bengio, Y., & Courville, A. (2016). Deep Learning. MIT Press.

Goodfellow lan, Bengio Yoshua, C. A. (2016). Deep Learning - Ian Goodfellow, Yoshua Bengio, Aaron Courville - Google Books. In MIT Press.

Grimes, W. N., Aytürk, D. G., Hoon, M., Yoshimatsu, T., Gamlin, C., Carrera, D., Nath, A., Nadal-Nicolás, F. M., Ahlquist, R. M., Sabnis, A., Berson, D. M., Diamond, J. S., Wong, R. O., Cepko, C., & Rieke, F. (2021). A High-Density Narrow-Field Inhibitory Retinal Interneuron with Direct Coupling to Müller Glia. Journal of Neuroscience, 41(28), 6018–6037. https://doi.org/10.1523/JNEUROSCI.0199-20.2021

Gu, Y., Chen, A., Zhang, X., Fan, C., Li, K., & Shen, J. (2021). Deep Learning based Cell Classification in Imaging Flow Cytometer. ASP Transactions on Pattern Recognition and Intelligent Systems, 1(2), 18–27. https://doi.org/10.52810/TPRIS.2021.100050

He, C., Liu, J., Zhu, Y., & Du, W. (2021). Data augmentation for deep neural networks model in EEGclassification task: A review. Front. Hum. Neurosci., 15, 765525.

Iqbal, M. S., Ahmad, I., Bin, L., Khan, S., & Rodrigues, J. J. P. C. (2021). Deep learning recognition of diseased and normal cell representation. Transactions on Emerging Telecommunications Technologies, 32(7), e4017. https://doi.org/10.1002/ett.4017

Jusuf, P. R., & Harris, W. A. (2009). Ptf1a is expressed transiently in all types of amacrine cells in the embryonic zebrafish retina. Neural Development, 4(1), 34. https://doi.org/10.1186/1749-8104-4-34

Ker, J., Bai, Y., Lee, H. Y., Rao, J., & Wang, L. (2019). Automated brain histology classification using machine learning. Journal of Clinical Neuroscience, 66, 239–245. https://doi.org/10.1016/j.jocn.2019.05.019

Khan, S. M., Liu, X., Nath, S., Korot, E., Faes, L., Wagner, S. K., Keane, P. A., Sebire, N. J., Burton, M. J., & Denniston, A. K. (2021). A global review of publicly available datasets forophthalmological imaging: barriers to access, usability, andgeneralisability. Lancet Digit. Health, 3(1), e51–e66.

Krieger, B., Qiao, M., Rousso, D. L., Sanes, J. R., & Meister, M. (2017). Four alpha ganglion cell types in mouse retina: Function, structure, and molecular signatures. PLOS ONE, 12(7), e0180091.. https://doi.org/10.1371/journal.pone.0180091

Krishnamoorthy, V., Jain, V., Cherukuri, P., Baloni, S., & Dhingra, N. K. (2008). Intravitreal Injection of Fluorochrome-Conjugated Peanut Agglutinin Results in Specific and Reversible Labeling of Mammalian Cones In Vivo. Investigative Ophthalmology & Visual Science, 49(6), 2643–2650. https://doi.org/10.1167/iovs.07-1471

Liu, X., Faes, L., Kale, A. U., Wagner, S. K., Fu, D. J., Bruynseels, A., Mahendiran, T., Moraes, G., Shamdas Mohithand Kern, C., Ledsam, J. R., Schmid Martin Kand Balaskas, K., Topol, E. J., Bachmann, L. M., Keane, P. A., & Denniston, A. K. (2019). A comparison of deep learning performance against health-careprofessionals in detecting diseases from medical imaging: asystematic review and meta-analysis. Lancet Digit. Health, 1(6), e271–e297.

Masland, R. H. (2001). The fundamental plan of the retina. Nature Neuroscience, 4(9), 877–886. https://doi.org/10.1038/nn0901-877

Noorbakhsh, J., Farahmand, S., Foroughi pour, A., Namburi, S., Caruana, D., Rimm, D., Soltanieh-ha, M., Zarringhalam, K., & Chuang, J. H. (2020). Deep learning-based cross-classifications reveal conserved spatial behaviors within tumor histological images. Nature Communications, 11(1), 6367. https://doi.org/10.1038/s41467-020-20030-5

Oei Ronald Wihal and Hou, G. (2019). Convolutional neural network for cell classification using microscope images of intracellular actin networks. PLOS ONE, 14(3), 1–13. https://doi.org/10.1371/journal.pone.0213626

Poché, R. A., & Reese, B. E. (2009). Retinal horizontal cells: challenging paradigms of neuraldevelopment and cancer biology. Development, 136(13), 2141–2151.

Ren, J. L., Yu, Q. X., Liang, W. C., Leung, P. Y., Ng, T. K., Chu, W. K., Pang, C. P., & Chan, S. O. (2018). Green tea extract attenuates LPS-induced retinal inflammation in rats. Scientific Reports, 8(1), 429. https://doi.org/10.1038/s41598-017-18888-5

Saha, S., Nassisi, M., Wang, M., Lindenberg, S., kanagasingam, Y., Sadda, S., & Hu, Z. J. (2019). Automated detection and classification of early AMD biomarkers using deep learning. Scientific Reports, 9(1), 10990. https://doi.org/10.1038/s41598-019-47390-3

Sandler, M., Howard, A., Zhu, M., Zhmoginov, A., & Chen, L.-C. (2018). MobileNetV2: Inverted Residuals and Linear Bottlenecks. 2018 IEEE/CVF Conference on Computer Vision and Pattern Recognition, 4510–4520. https://doi.org/10.1109/CVPR.2018.00474

Shibata, M., Okamura, K., Yura, K., & Umezawa, A. (2020). High-precision multiclass cell classification by supervisedmachine learning on lectin microarray data. Regen. Ther., 15, 195–201.

Shifat-E-Rabbi, M., Yin, X., Fitzgerald, C. E., & Rohde, G. K. (2020). Cell image classification: A comparative overview. Cytometry A, 97(4), 347–362.

Shorten, C., & Khoshgoftaar, T. M. (2019). A survey on Image Data Augmentation for Deep Learning. Journal of Big Data, 6(1), 60. https://doi.org/10.1186/s40537-019-0197-0

Strettoi, E., Mears, A. J., & Swaroop, A. (2004). Recruitment of the Rod Pathway by Cones in the Absence of Rods. The Journal of Neuroscience, 24(34), 7576. https://doi.org/10.1523/JNEUROSCI.2245-04.2004

Tabibu, S., Vinod, P. K., & Jawahar, C. V. (2019). Pan-Renal Cell Carcinoma classification and survival prediction from histopathology images using deep learning. Scientific Reports, 9(1), 10509. https://doi.org/10.1038/s41598-019-46718-3

Ting, D. S. W., Pasquale, L. R., Peng, L., Campbell, J. P., Lee, A. Y., Raman, R., Tan, G. S. W., Schmetterer, L., Keane, P. A., & Wong, T. Y. (2019). Artificial intelligence and deep learning in ophthalmology. Br. J. Ophthalmol., 103(2), 167–175.

Wan, J., & Goldman, D. (2017). Opposing actions of Fgf8a on Notch signaling distinguish twoMuller glial cell populations that contribute to retina growthand regeneration. Cell Rep., 19(4), 849–862.

Wilson, A. M., & Di Polo, A. (2012). Gene therapy for retinal ganglion cell neuroprotection inglaucoma. Gene Ther., 19(2), 127–136.

Zele, A. J., & Cao, D. (2014). Vision under mesopic and scotopic illumination. Front. Psychol., 5, 1594.

Zhang, L., Lu, L., Nogues, I., Summers, R. M., Liu, S., & Yao, J. (2017). DeepPap: Deep Convolutional Networks for Cervical Cell Classification. IEEE Journal of Biomedical and Health Informatics, 21(6), 1633–1643. https://doi.org/10.1109/JBHI.2017.2705583

